# Genomic profiling of T cell activation reveals dependency of memory T cells on CD28 costimulation

**DOI:** 10.1101/421099

**Authors:** Dafni A. Glinos, Blagoje Soskic, Luke Jostins, David M. Sansom, Gosia Trynka

**Affiliations:** Wellcome Sanger Institute, Wellcome Trust Genome Campus, Hinxton CB10 1SA, UK.; Kennedy Institute of Rheumatology, University of Oxford, Roosevelt Drive, Oxford OX3 7FY, UK.; Big Data Institute, University of Oxford, Roosevelt Drive, Oxford OX3 7FZ, UK.; Christ Church, St. Aldates, Oxford OX1 1DP, UK.; UCL Institute of Immunity and Transplantation, Royal Free Hospital, London NW3 2PF, UK.

## Abstract

T cell activation is a critical driver of immune response and if uncontrolled, it can result in failure to respond to infection or in excessive inflammation and autoimmunity. CD28 costimulatory pathway is an essential regulator of CD4 T cell responses. To deconvolute how T cell receptor (TCR) and CD28 orchestrate activation of human CD4 T cells we stimulated cells using varying intensities of TCR and CD28 signals followed by gene expression profiling. We demonstrate that T-helper differentiation and cytokine expression are controlled by CD28. Strikingly, cell cycle and cell division are sensitive to CD28 in memory cells, but under TCR control in naive cells, in contrast to the paradigm that memory cells are CD28-independent. Using a combination of chromatin accessibility and enhancer profiling, we observe that IRFs and Blimp-1 (PRDM1) motifs are enriched in naive and memory T cells in response to TCR. In contrast, memory cells initiate AP1 transcriptional regulation only when both TCR and CD28 are engaged, implicating CD28 as an amplifier of transcriptional programmes in memory cells. Lastly, we show that CD28-sensitive genes are enriched in autoimmune disease loci, pointing towards the role of memory cells and the regulation of T cell activation through CD28 in autoimmune disease development. This study provides important insights into the differential role of CD28 in naive and memory T cell responses and offers a new platform for design and interpretation of costimulatory based therapies.

**One-sentence summary:** Genomic profiling of CD4 T cell activation reveals a sensitivity switch from TCR in naive to CD28 in memory cells.

## Introduction

The ability of T cells to respond to pathogens whilst remaining tolerant to host antigens is critical for human health. T cell stimulation generally occurs in secondary lymphoid tissues where T cells interact with professional antigen presenting cells (APCs). Here, two coordinated signals are delivered: the first via T cell receptor (TCR) recognising antigen bound to MHC molecules and the second provided by APCs via upregulation of costimulatory ligands. In this regard, CD28 is the main costimulatory receptor expressed by T cells which interacts with CD80 and CD86 ligands on APCs. The coordination of TCR and CD28 signals is essential for T cell activation, proliferation, differentiation and survival, making the CD28 pathway a key checkpoint for controlling T cell responses (Rowshanravan et al., 2018).

Nevertheless, it remains controversial whether CD28 engagement differentially impacts activation of naive and memory T cells. Previous studies have suggested that memory T cells have lower costimulation thresholds or conversely, that naive cells have a greater requirement for costimulatory signals (Croft et al., 1994; Dubey et al., 1995; London et al., 2000; Luqman and Bottomly, 1992). This has given rise to a widely perceived notion that, in contrast to naive T cells, previously activated or memory T cells do not require CD28 costimulation. However, there is evidence that this may not be the case (van der Heide and Homann, 2016; Ville et al., 2015) and that CD28 costimulation is important to numerous aspects of functional competence for memory T cells (Borowski et al., 2007; Fröhlich et al., 2016; Linterman et al., 2014; Ndlovu et al., 2014). These contrasting conclusions are likely influenced by the experimental systems used, including the nature and intensity of the TCR signal. A further important issue is how CD28 responsiveness is measured. For example, the requirement for CD28 to induce T cell proliferation appears lower than that for T follicular helper (Tfh) differentiation (Wang et al., 2015), a process essential for the regulation of antibody production by B cells. As such, different T cell processes may vary in their degree of dependence on CD28 costimulation.

The level of CD28 costimulation vary considerably in different immunological settings. For example, the presence of regulatory T cells (Tregs) expressing CTLA-4, which degrades CD80 and CD86 ligands (Qureshi et al., 2011) will influence CD28 costimulation. Indeed, deficiency in expression of CTLA-4 is associated with development of profound autoimmune diseases (Kuehn et al., 2014; Lo et al., 2015; Schubert et al., 2014; Tivol et al., 1995) due to increased CD28 signalling (Tai et al., 2007; Tivol et al., 1997). In addition, severe adverse reactions were observed during trials of a CD28 agonistic antibody TGN1412, which triggered a cytokine storm from effector memory T cells (Eastwood et al., 2010; Hünig, 2012). Furthermore, excessive activation of memory T cells is also a hallmark of many common complex immune diseases, such as autoimmune arthritis and systemic lupus erythematosus (SLE) (Haufe et al., 2011; Kshirsagar et al., 2013). Genome-wide association studies (GWAS) have mapped numerous risk variants to loci encoding genes in T cell stimulatory pathways, including the *CD28*, *CTLA4* and *ICOS* genes located at 2q33.2 (Dubois et al., 2010; Fortune et al., 2015; Okada et al., 2014; Onengut-Gumuscu et al., 2015). While the exact effects of the associated variants are unknown, their mapping to the non-coding regions of the genome suggests effects on regulation of gene expression (Fairfax et al., 2014; Westra et al., 2013), which implies modulation of activity of costimulatory pathways in biology of immune-mediated diseases. Therefore, understanding how varying levels of TCR and CD28 costimulation impact gene expression and the ensuing T cell response, has significant implications in understanding susceptibility to diseases, in particular autoimmunity and cancer.

To understand the requirement of TCR and CD28 in the activation of both naive and memory human CD4 T cells we stimulated cells with varying intensities of TCR and CD28 signals. We used gene expression and chromatin activity profiling to map transcriptional processes underlying T cell activation. We show that the major effector cell functions, such as T helper (Th) differentiation, expression of chemokine receptors and cytokines, are strongly influenced by CD28 in both naive and memory T cells. Strikingly, cell division is markedly differently controlled between the two cells, which we find to be controlled by CD28 in memory cells whilst predominantly driven by TCR in naive cells. Finally, we show that loci associated to common immune diseases are enriched in CD28-sensitive genes, pointing towards the important role of CD28 costimulatory pathway in disease pathogenesis. Taken together, this study provides key insights into the relative impact of TCR and CD28 in the activation of human naive and memory T cells.

## Results

### Experimental approach

To study the influence of varying TCR and CD28 signals we sorted human blood CD4+ CD25-T cells and stimulated them with high or low concentrations of soluble anti-CD3 (an antibody against the signalling component of the TCR complex; referred to as high and low TCR) and CHO cells expressing CD86, the CD28 ligand (referred to as high and low CD28) (**Fig. 1A** and **Materials and Methods**). We have previously shown that on their own CHO cells do not activate T cells (30). Indeed, in our experimental setup we used resting cells where T cells were cultured with fixed CHO cells and we observed no cell activation, as shown by the lack of CD25 upregulation (**Suppl. Figure S1**). The advantage of using CHO-CD86 over other stimulation systems is that it delivers a natural CD28 ligand, which was documented to be the main CD28 ligand in vivo (Borriello et al., 1997). We also wanted to deconvolute gene expression programmes controlled individually by TCR and CD28. On its own, CD86 is insufficient to deliver strong enough signal for CD28 to activate cells. However, this can be achieved with anti-CD28 antibody. To maintain consistent cell culture conditions we stimulated T cells with individual anti-CD3 and anti-CD28 antibodies in the presence of CHO cells expressing Fc-gamma Receptor II (CHO-FcR) to provide crosslinking, rather than immobilizing antibodies on the plate. Following a 16h stimulation we sorted activated CD25+ cells into naive (CD45RA+) and memory (CD45RA-) subsets (**Materials and Methods**). To map the transcriptional programmes underlying TCR and CD28 sensitivity in naive and memory T cells, we profiled gene expression with RNA-seq, chromatin accessibility with ATAC-seq and active enhancers and promoters with H3K27ac ChIPmentation-seq (Schmidl et al., 2015). As expected, across different stimulatory conditions we observed a variable proportion of activated T cells (11-78% of all cells) (**Suppl. Figure S1**). However, by sorting only activated T cells we ensured that the measured gene expression reflected cell activation state induced by different signal intensities, while not being confounded by the variable percentage of cells that had undergone activation.

**Figure 1:**
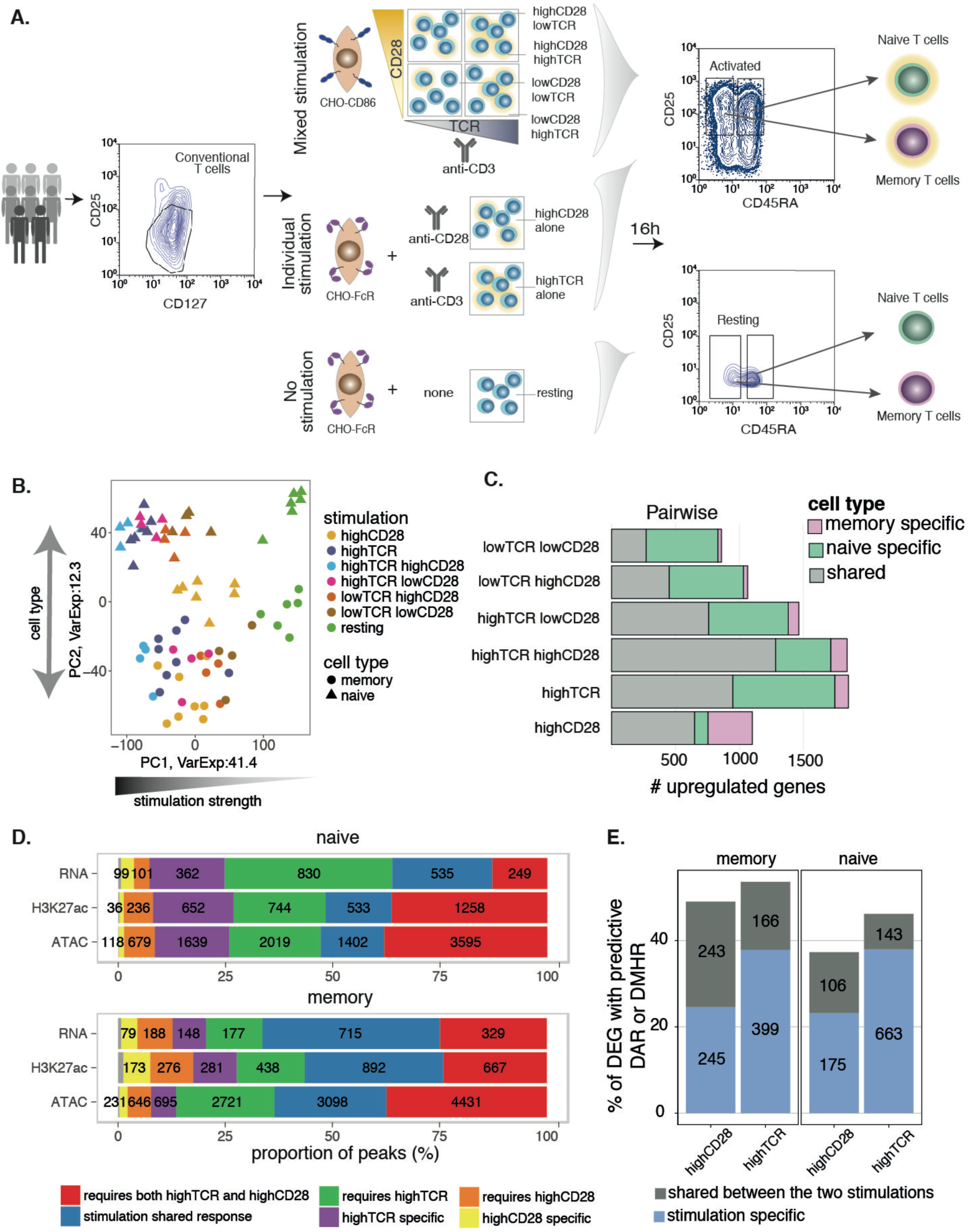
**A**. Overview of the study design. CD4 T cells were isolated from eight healthy individuals and cultured in six different stimulatory conditions, which included variable concentrations of anti-CD3 and anti-CD28 stimuli. In parallel, resting cells were cultured as a control. To ensure we measured cellular response to successful stimulation, we generated sequencing data from sorted activated naive and memory cells identified as CD4+CD25+CD45RA^high^ and CD4+CD25+CD45RA^low^, respectively. **B.** Principal component analysis using the expression of all genes. The first two components explain collectively 53.7% of the observed variability and correlate with stimulation strength and cell type. Each dot corresponds to an individual sample, colored by stimulation and shaped according to the cell type. **C**. Number of upregulated genes upon stimulation defined by pairwise differential expression test between stimulated cells and resting cells (fold-change ≥ 2 and FDR ≤ 0.05). **D**. Proportion of differentially expressed genes (DEG), differentially accessible regions (DARs) and differentially modified histone regions (DMHRs) upon stimulation with both stimuli, strong TCR alone (high TCR) or strong CD28 alone (high CD28). Written inside the barplot is the corresponding number of detected differential changes. The coloring represents different stimulatory conditions. Grey indicates a small number of peaks that were shared between high TCR alone and high CD28 alone stimulations. **E**. Proportion of differentially expressed genes that have at least one differentially regulated region nearby (< 150kb away from TSS) as detected by ATAC-seq (DAR) and H3K27ac ChM-seq (DMHR).

### Naive and memory T cells operate different gene expression programmes upon activation

We first applied principal component analysis (PCA) to the RNA-seq data and observed that PC1 reflected cell stimulation while PC2 corresponded to cell type (**Fig. 1B and Suppl. Fig. S2A**). Indeed, the majority of the gene expression variance was explained by stimulation (47%) and by the different cell type (10%) (**Suppl. Fig. S2B** and **Materials and Methods**). The separation of naive and memory T cells by PC2 indicated clear differences in transcriptional responses between naive and memory T cells. The different conditions clustered together, capturing the gradient of stimulation intensity in both cell types, with high levels of both TCR and CD28 stimuli being the furthest separated from unstimulated cells on PC1 and intermediate intensity of stimuli mapping in between. Surprisingly, strong CD28 alone (FcR-anti-CD28) was amongst the lower responding conditions in naive cells yet it clustered with the more highly stimulated conditions in memory cells. Furthermore, naive cells stimulated with strong CD28 alone clustered towards the memory cells along PC2. Thus, the transcriptional program of cell activation is modulated by the intensity of TCR and CD28 signals, and CD28 stimulation enriches for the characteristics of memory T cells.

To understand cell type specific responses induced by the different stimuli, we compared gene expression profiles between resting and stimulated naive and memory T cells (false discovery rate (FDR) ≤ 0.05 and fold-change ≥2; **Fig. 1C** and **Suppl. Table S1**). As expected, the majority of the upregulated genes were shared between the two cell types, however, naive cells displayed a larger number of differentially upregulated genes (DEGs) (average of 1,240 genes) than memory cells (average of 840 genes), with an exception of CD28 stimulation alone, where more genes were upregulated in memory cells. This likely reflects the greater changes in gene expression levels resulting from transitioning to activation from a deeper quiescent state in naive cells. Indeed, differential expression analysis between naive and memory cells in the resting state revealed a larger number of genes expressed highly in memory cells compared to naive (**Suppl. Fig. S3**).

When looking globally at the whole genome maps of H3K27ac and chromatin accessible regions (ATAC-seq), memory cells were characterised by more peaks in the resting state in both ATAC-seq (17.5% more peaks) and H3K27ac ChM-seq (10.6% more peaks). In order to gain a better understanding of the dynamic responses upon stimulation, we performed the same comparisons as with the RNA-seq data. At the mRNA level we observed 35% of the genes to be differentially expressed upon stimulation, however the chromatin regulatory landscape showed far fewer changes; only 8% of the chromatin changed in accessibility and 9% of the regions changed in histone acetylation. This is probably due to the fact that the majority of these genes are already expressed at low levels, which would be reflected by chromatin being already open and enhancers being marked by H3K27ac acetylation.Furthermore, similarly to the differential gene expression, the majority of differential histone modified regions (DMHRs; 54%; see **Materials and Methods**) and differentially accessible regions (DARs; 58%) were shared across responses to TCR, CD28, or both. However, as with the gene expression analysis, we observed a larger proportion of acetylation changes driven by CD28 alone in memory compared to naive cells (Fisher’s exact test RNA p-value = 4.9×10^−11^; H3K27ac p-value < 2.2×10^−16^), indicating that CD28, in line with our observations for the RNA, is also more potent in inducing chromatin changes in memory T cells. Conversely, in response to strong TCR stimulation alone, we detected a larger proportion of differentially acetylated H3K27 regions and chromatin accessible sites in naive cells compared to memory cells (Fisher’s exact test RNA p-value < 2.2×10^−16^; H3K27ac p-value < 2.2×10^−16^; ATAC p-value < 2.2×10^−16^; **Fig. 1D**). As such, the observed high number of stimulus-specific differentially regulated regions suggests that there are unique chromatin remodelling changes acquired in both a cell type and stimulus-specific context.

Given the observation that TCR and CD28 induced different number of changes in chromatin accessibility and histone modifications, we next assessed if there was a difference in the proportion of these chromatin activity changes that were also predictive of the observed differential gene expression (see **Materials and Methods**) (Wang et al., 2013). We found that DARs and DMHRs were predictive of a large proportion of upregulated genes (mean 46.5%; **Fig. 1E**). Although, globally, TCR induced more changes in chromatin activity, the percentage of upregulated genes predicted by DMHRs and DARs was similar between TCR and CD28 (**Suppl. Fig. S4**). We observed that the majority of genes for which we had assigned predictive differentially regulated regions included peaks differentially regulated upon a specific stimulus (naive TCR 82.3%; memory TCR 70.6%; naive CD28 62.23%; memory CD28 50.2%). This indicates that each stimulus uniquely contributes to the gene regulatory landscape of a cell.

### T cell effector functions are predominantly controlled by CD28

Based on the above observations we sought to better understand the role of each stimulatory signal in gene upregulation in naive and memory T cells. To classify genes as either CD28-or TCR-sensitive, we used two models (linear and switch modes) of gene expression where the linear model reflected changes in stimulation intensity whereas the switch model reflected a digital on/off state (**Fig. 2A**). In both cell types together, we were able to assign a unique stimulus sensitivity to 1,567 genes, meaning that the expression of these genes was sensitive to either TCR or CD28 intensity. We observed that the majority of stimulus-sensitive genes (88%) followed the linear model (**Suppl. Fig. S5A**).

**Figure 2:**
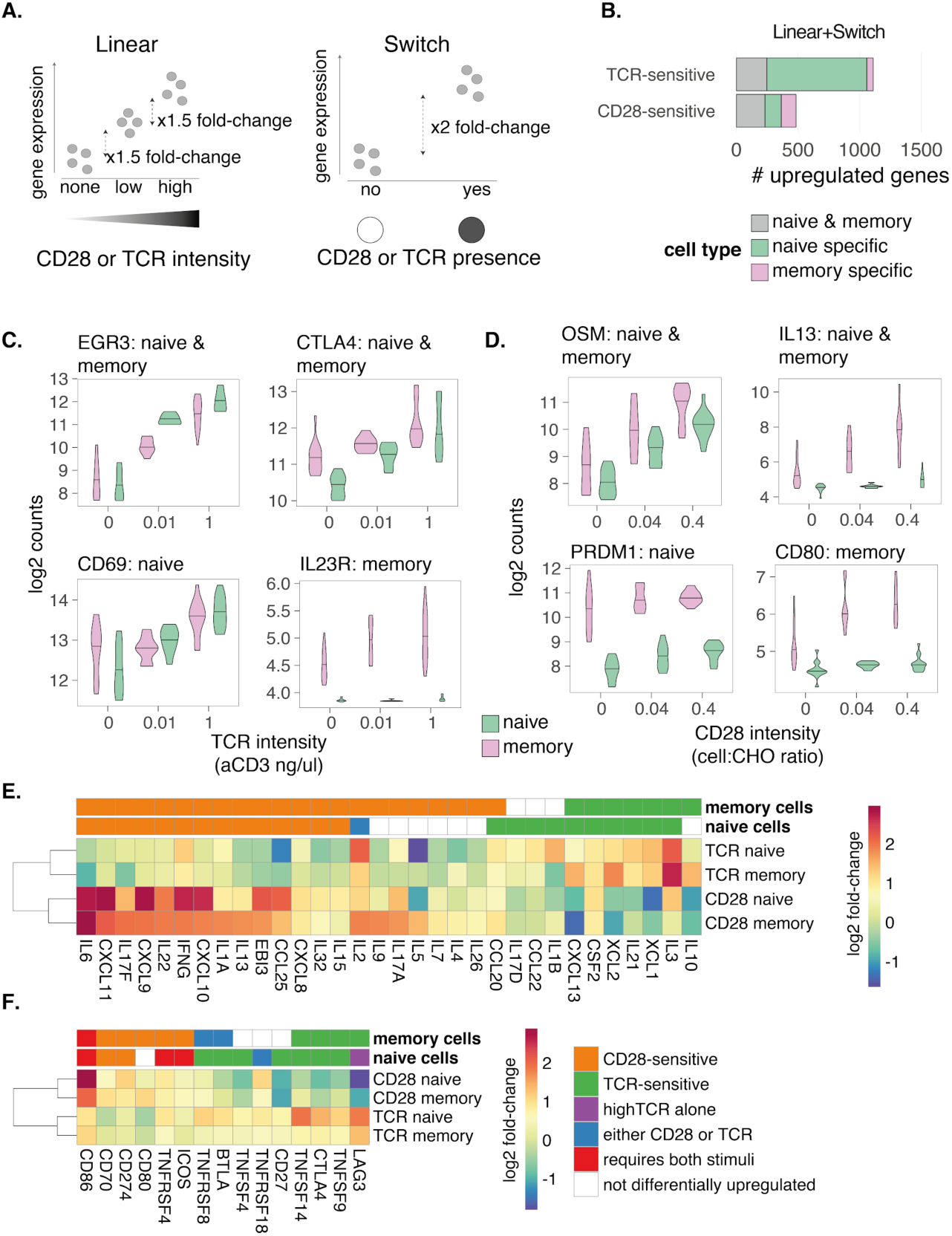
**A**. Classification of genes as CD28-or TCR-sensitive using different models of gene expression. In the linear model, we required a linear increase of gene expression along with stimulus intensity (incremental fold-change ≥ 1.5 in gene expression), separately evaluating naive and memory cells. Genes that did not follow the linear model were tested for the switch model. Here, we assumed an “on-and-off” mode of expression where a gene is significantly upregulated (fold-change ≥ 2) in response to the presence of either CD28 or TCR. In both of these models, we used all seven conditions, e.g. when testing for CD28-sensitive genes we grouped the TCR alone stimulation with the resting, since neither received CD28 signal. A gene was classified in one of the two categories without overlap and prioritised for the linear model. **B**. Comparison of the number of genes in naive and in memory cells under TCR control or CD28. **C+D**. Selected examples of TCR-or CD28-sensitive genes. The x-axis corresponds (**C**) to the level of TCR (anti-CD3 antibody in μg/ml) or (**D**) to the level of CD28 (proportion of T cells to CHO-CD86 cells) and the y-axis corresponds to the log2 counts of gene expression. Labels next to the gene name indicate the cell type in which stimulus sensitivity is observed. **E.** Cytokines and chemokines, and **F**. costimulatory ligands and receptors that are sensitive to at least one stimulus in at least one cell type. Colouring represents the log2 fold change of gene expression.

We next assessed if naive and memory cells differed in sensitivity to the two stimuli. We observed that most of the stimulus-sensitive genes in naive T cells were TCR-sensitive (1,057; **Fig. 2B** and **Suppl. Table S2**), whereas a smaller number of genes was CD28-sensitive (n=363). Conversely, in memory cells we observed that a larger number of genes was CD28-sensitive (n=351) than TCR-sensitive (n=299). As such, a both TCR sensitive genes and CD28 sensitive genes were unevenly distributed between naive and memory cells, with a shift towards naive cells for TCR genes (Fisher’s exact test p-value < 2.2×10^−16^) and a shift towards memory cells for CD28 genes (Fisher’s exact test p-value < 2.14×10^−5^). Based on the pairwise comparisons across the six conditions against the resting state, we defined a group of genes that was upregulated upon stimulation. Among this group we observed that the expression of 1,228 genes in naive cells (55%) and only 490 genes in memory cells (29%) was sensitive to a single stimulus (**Suppl. Fig. S5B**). This indicated that the majority of the upregulated genes in memory cells either responded to both stimuli (i.e. TCR or CD28 were both capable of driving the response) or they were truly CD28 costimulation dependent, requiring TCR and CD28 together.

Since the majority of human T cell stimulation experiments use both TCR and CD28 to activate cells, it is unclear which T cell functions are controlled by TCR and which by CD28, and how they differ between naive and memory cells. Using this approach, we found that many classical T cell activation markers such as *EGR2*, *EGR3* and *CTLA4* were TCR-sensitive in both cell types, while *TNFRSF8* (encodes for CD30) and *CD69* were TCR-sensitive in naive cells only (**Fig. 2C**), possibly explained by the fact that they were already expressed at high levels in resting memory cells (*TNFRSF8* log2FC = 1.19 and *CD69* log2FC = 1 between resting memory and naive cells) and thus not detected as differentially expressed upon memory T cell stimulation (**Suppl. Fig. S3**). In contrast, we found that the majority of cytokines and chemokines were CD28-sensitive (**Fig. 2E** and **Suppl. Fig. S5C**). Among the chemokines, we observed that *CXCL9*, *CXCL10*, *CXCL11* and the *CCL25* chemokine were CD28-sensitive in both cell types. Notably, the expression of cytokines essential for the differentiation of the major Th subsets were also found to be under CD28 control, including *IFNG* (Th1), *IL4* and *IL13* (Th2) and *IL17A*, *IL17F* and *IL22* (Th17). In addition, we observed that expression of the Treg transcription factor, *FOXP3,* was also dependent on CD28 stimulation. Taken together, these results demonstrate that a number of genes associated with effector functions of CD4 T cells are predominantly controlled by the CD28 pathway in both naive and memory cells.

Different co-stimulatory and co-inhibitory receptors along with their ligands function at different stages of the T cell differentiation cascade which might explain a differential requirement on TCR or CD28 for their expression (Chen and Flies, 2013). We therefore also investigated the control of expression of costimulatory ligands and receptors, since they play a key role in T cell activation (**Fig. 2F** and **Suppl. Fig. S5D**). The expression of *CD28* itself was only upregulated upon strong TCR stimulation alone, suggesting that TCR signalling makes cells more receptive to CD28 engagement. However, the presence of CD28 stimulus impeded an increase in its expression, highlighting that this is not a self-reinforcing process. Other co-stimulatory/co-inhibitory receptors involved in the co-regulation of T cells were CD28-sensitive, in both naive and memory T cells. These included, the *CD27*-*CD70* costimulatory pair and *CD274* (PD-L1) (the expression of *PD1* was not detectable). On the other hand, *CD80* and *ICOS* were CD28-sensitive specifically in memory cells. Thus, we were able to detect different modes of regulation for *CD28*, *CTLA4* and *ICOS*, despite the fact that they are all encoded within the same 260kb locus on chromosome 2. This implies complex mechanisms of gene expression regulation in the two cell types and a fine-tuned control highlighting that co-located genes can be regulated by different modes of stimulation.

### DNA replication and proliferation are driven by different stimuli in naive and memory T cells

To determine whether genes sensitive to CD28 or TCR regulate the same cellular processes in naive and memory T cells we tested if these genes were over-represented in hallmark functional pathways (Liberzon et al., 2015) (**Materials and Methods**; **Fig. 3A** and **Suppl.Fig. S4**). We observed seven shared pathways enriched in both cell types. For example, *IL2 signalling through STAT5* and *TNF-α signalling via NF-κβ* were significantly enriched in both naive and memory cells and sensitive to both TCR and CD28. However, the expression of the majority of genes broadly classified as immune cell pathways, such as *IL6 signalling through the Jak/Stat3* and *interferon α and ɣ response*, were CD28-sensitive in both cell types. In contrast, genes that are targets of Myc and E2F transcription factors were controlled by TCR in naive cells, which is concordant with *MYC* and *E2F6* genes being TCR-sensitive, and also with previous studies in mice (Allison et al., 2016).

**Figure 3:**
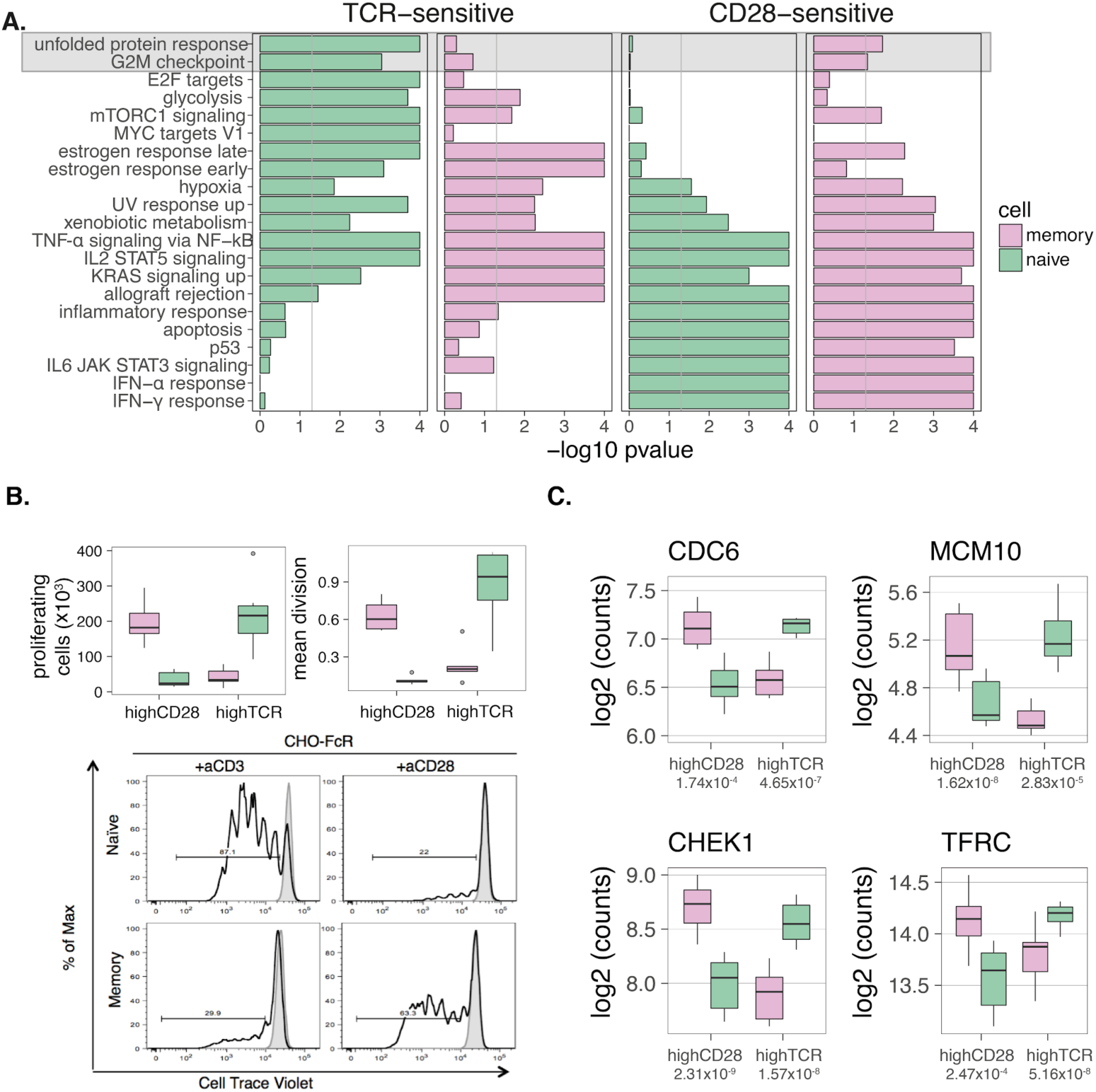
**A.** Hallmark pathways enriched for TCR-sensitive and CD28-sensitive genes in naive and memory cells. **B.** CTV labelled naive and memory T cells were stimulated with anti-CD3 or anti-CD28 in the presence of CHO-FcR. The proliferation of naive and memory T cells was measured by flow cytometry five days following stimulation. Upper panel boxplots show the number of proliferating cells. Lower panel flow cytometry histograms show proliferation of naive and memory T cells measured by dilution of CTV. **C.** Gene expression profiles of selected switcher genes, genes that change stimulus sensitivity between the two cell types. The number below the stimulus name represents the significance of the difference between naive and memory T cells derived using t-test.

Interestingly, we observed that some pathways were differentially sensitive to TCR and CD28 in the two cell types. Most notably, the *G2M checkpoint*, which marks DNA replication and cell division, was CD28-sensitive in memory cells but TCR-sensitive in naive cells, suggesting that commitment to cell division is more dependent on TCR in naive cells but driven by CD28 in memory cells. To functionally test this observation we stimulated CellTrace Violet (CTV) labelled naive and memory T cells with either anti-CD3 or anti-CD28 crosslinked by CHO cells expressing an FcR (**Fig. 3B**). Five days following stimulation, T cell division was measured by flow cytometry. In accordance with the results from the gene expression, naive T cells proliferated extensively following TCR crosslinking but mounted poor responses to CD28. In contrast, cross-linking CD28 was sufficient to induce division in memory cells whereas TCR stimulation was clearly less effective. Thus, although combined TCR and CD28 costimulation is generally utilised to trigger T cell proliferation, our data indicates a division of labour between these stimuli, where control of cell cycle in naive cells is generally TCR-sensitive, whilst in memory cells it is more dependent on CD28.

We noticed that there were three genes (*CDC6*, *CDC20* and *CHEK1*) driving the enrichment of the G2M pathway, which were TCR sensitive in naive cells but switched to CD28 sensitivity in memory cells. We therefore sought to identify if there were more “switching” genes present in our dataset, i.e. genes sensitive to a different stimulus between the two cell types. We identified a group of 18 genes that were TCR-sensitive in naive cells and changed to CD28 sensitivity in memory cells. Among others, we identified transferrin receptor *TFRC* (CD71), which is necessary for iron uptake and fueling of the proliferation, as well as *MCM10* that is an important factor initiating DNA helicase activity and replication (**Fig. 3C** and **Suppl. Table S3**). Together these data highlight the enrichment of cell cycle/DNA replication pathways as targets for CD28 in memory T cells which, in contrast, are controlled by the TCR in naive cells.

### AP1 initiated transcriptional cascade is costimulation dependent in memory cells

To understand how TCR and CD28 stimuli exert effects on gene expression we tested for enrichment of transcription factor binding sites (TFBSs) in DARs identified by ATAC-seq that were concordant with gene expression (**Materials and Methods**, **Fig. 4A**). We tested 242 transcription factors with detectable levels of gene expression in our dataset.

**Figure 4:**
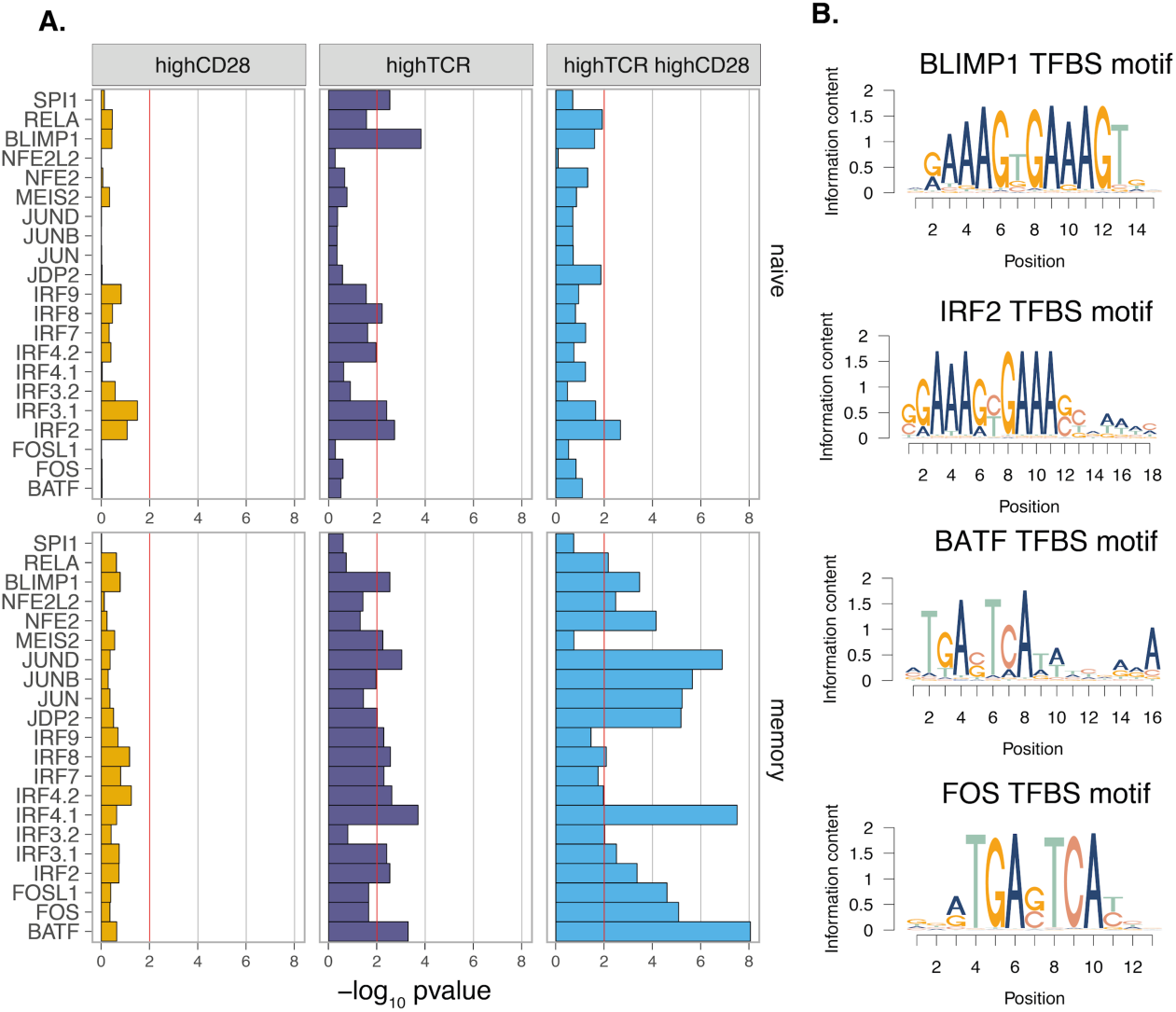
**A**. Transcription factor binding sites (TFBS) enriched in the differentially regulated ATAC-seq peaks that were assigned to predict gene expression changes. **B**. Position weight matrices (PWMs) for the enriched TFBS motifs.

We identified 5 different motifs enriched in naive cells and 20 motifs in memory cells (p-value ≤ 0.01). In naive T cells we observed an enrichment for Blimp-1, a transcriptional repressor that maintains T cell homeostasis (Martins et al., 2006), combined with interferon response elements, IRF2, IRF3 and IRF8. Increases in TCR signalling have been shown to induce the IRF4 transcription factor, which mediates Blimp-1 abundance in mice (Man et al., 2013). As expected, the enrichment was driven by TCR, with little enhancement in response to CD28 costimulation. Blimp-1 and the identified IRFs recognize a very similar motif (**Fig.4B**), and antagonize each others binding *in vitro* (Doody et al., 2010).

In memory T cells we observed that the profile driven by TCR was similar to naive cells, consisting of a combination of Blimp-1 with interferon response elements. However, in marked contrast, a much more robust response was observed in the presence of CD28 co-stimulation. Notably, this was characterised by the enrichment of AP1 transcription factors (**Fig. 4A**). Specifically, only IRF4, JunD and BATF transcription factor binding sites co-occurred in regions that changed in response to strong TCR alone, whereas c-Fos, FOSL1, c-jun, jun-B and JDP2 all required the presence of CD28. C-jun and c-Fos constitute the backbone of AP1, a transcription factor that plays an important role in the induction of the immune response. Thus our observations are consistent with previous work which suggested that CD28 regulates the expression and activity of AP1 transcription factors (Edmead et al., 1996; Fraser et al., 1991; Shapiro et al., 1997). Interestingly, BATF and c-jun transcription factors can also form a heterodimer and cooperate with IRF4 and recognise AP1–IRF composite elements (AICEs) in pre-activated CD4 T cells (**Fig. 4B**) (Li et al., 2012). These transcription factors play a crucial role in the initiation of transcriptional programs specific to T cell activation and cell division.

### Immune GWAS loci are enriched for CD28-sensitive genes

The role of T cell activation in the development of immune-mediated diseases is well established and SNPs nearby genes relevant to T cell activation, differentiation and trafficking have been implicated through GWAS (Farh et al., 2015; Hu et al., 2014; Okada et al., 2014; Trynka et al., 2013). We sought to investigate if immune disease associated loci were enriched for the genes we identified as TCR or CD28-sensitive, thereby implicating the involvement of either of these stimulatory pathways in disease pathogenesis.

In our enrichment analysis (see **Materials and Methods** and **Suppl. Table S4**) we tested eight immune-mediated conditions, Crohn’s disease (CD), ulcerative colitis (UC), celiac disease (CeD), rheumatoid arthritis (RA), type-1 diabetes (T1D), systemic sclerosis (SSc), multiple sclerosis (MS) and psoriasis (PSO). We used bone mineral density (BMD) as a negative control as we would not expect to observe significant enrichment among BMD loci. The majority of the tested immune diseases were more strongly enriched (permuted p-value < 0.01) for CD28-sensitive than TCR-sensitive genes. One exception was T1D where genes sensitive to TCR showed a higher enrichment (**Fig. 5A**). In addition, we observed that TCR-sensitive genes were enriched in CeD loci in both cell types (permuted p-value = 0.0097) and in RA (permuted p-value = 0.0096) and MS (permuted p-value = 0.0054) specifically in memory cells. Taken together, this indicates that the variants associated with common immune-mediated diseases could regulate the expression of genes sensitive to CD28 and through this pathway modulate the outcome of T cell activation.

**Figure 5.**
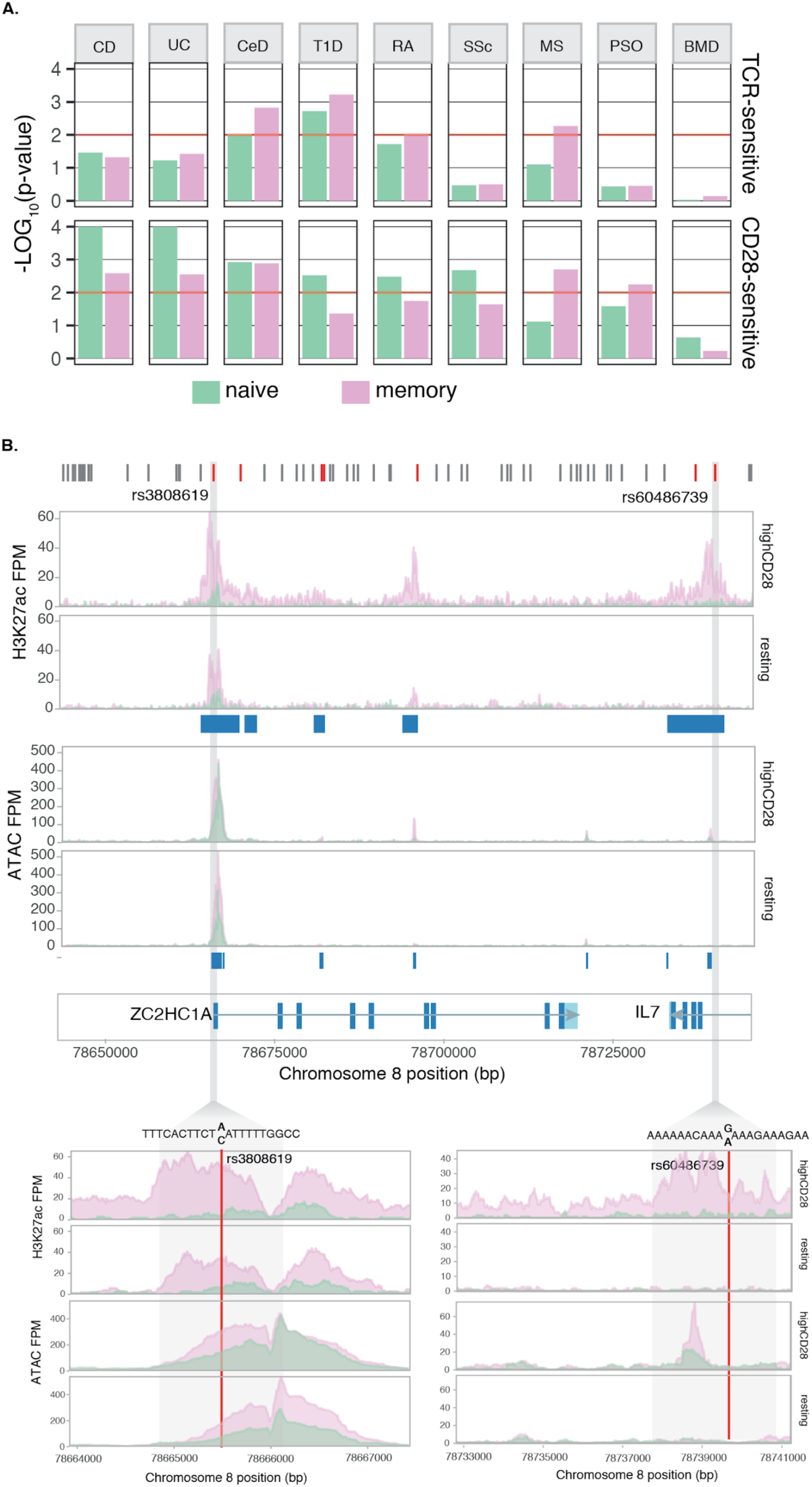
**A.** Enrichment of TCR-and CD28-sensitive genes in immune-mediated disease loci. Bone mineral density (BMD) is used as a negative control. (Crohn’s disease - CD, ulcerative colitis - UC, celiac disease - CeD, type-1 diabetes - T1D, rheumatoid arthritis - RA, systemic sclerosis - SSc, multiple sclerosis - MS, psoriasis - PSO). **B.** MS associated locus containing two genes, *ZC2HC1A* and *IL7,* that are CD28-sensitive. In the upper panel in red are indicated all the SNPs in LD with previously reported GWAS index variant, rs1021156. Of these, the two highlighted variants, rs3808619 and rs60486739, overlap CD28-upregulated H3K27ac peaks and are predicted to disrupt and IRF binding site.

The majority of immune disease associated genetic variants fall in the non-coding regions of the genome and previous studies showed that disease associated variants are enriched in active enhancers (Trynka et al., 2013, 2015). We therefore investigated if disease SNPs map within active or open chromatin regions as defined by H3K27ac or ATAC peaks near the genes driving the enrichment. Given that the majority of peaks were shared between the cell types and stimuli, these chromatin marks were uninformative in discriminating if disease SNPs were more significantly enriched in specific conditions. However, we observed that, on average across traits, 73% of the genes driving the enrichment also had at least one disease associated variant overlapping an active promoter or enhancer. We therefore tested whether any of these SNPs disrupted a TFBS, limiting our analysis to the TFs that we had previously identified as enriched. We found 13 unique SNPs that disrupted a binding motif (**Table 1**). Two SNPs, associated with MS and in high LD with the reported index variant rs1021156 (r^2^>0.8), disrupted the IRF TFBS within the *ZC2HC1A*/*IL7* locus (**Fig. 5B**). The first one, rs3808619 is localised in the promoter of *ZC2HC1A* and the risk allele led to decreased binding by IRF family of transcription factors (**Fig. 5B**). The second variant, rs60486739, is located in intron 3 of *IL7*, and the minor allele led to increased binding by IRF transcription factors (**Fig. 5B**). Both genes were sensitive to CD28 stimulation in memory cells and the H3K27ac peak that contained the rs60486739 variant was only present in stimulated memory cells, suggesting a potential functional role of the variant in modulating TF binding in this enhancer and affecting the expression levels of the gene.

**Table 1:** Transcription factor binding sites in open chromatin or H3K27ac peaks that are disrupted by immune-disease associated variants. **SNP** - disease SNP residing within an ATAC peak and disrupting TFBS; **All** - SNP alleles; **TF** - transcription factor recognising the binding site; **Delta PWM** - difference in PWM score between the reference and the alternative allele; **Trait** - disease for which the association was reported (T1D - type-1 diabetes, CeD - celiac disease, CD - Crohn’s disease, UC - ulcerative colitis, RA - rheumatoid arthritis, MS - multiple sclerosis, PSO - psoriasis, SSc - systemic sclerosis); **Cell** - cell type in which disease gene shows stimulus sensitivity; **Stim** - stimulus to which the gene is sensitive; **Gene** - gene that is stimulus-sensitive and maps to the disease locus.

| SNP | All | Gene | Stimulus | Cell | TF | PWM delta | Trait |
| --- | --- | --- | --- | --- | --- | --- | --- |
| rs2852151 | G/A | *SEH1L* | TCR | naive | RELA | -132 | T1D, CeD |
| rs3181365 | A/T | *TNFSF15* | CD28 | memory | BLIMP1 | -249 | CD, UC |
| rs6715826 | T/G | *MYO1B, STAT1* | CD28 | memory | BLIMP1 | -55 | CeD, RA, CD, UC, MS |
| rs2548530 | T/C | *ERAP2* | CD28 | naive | IRFs | -159 | CD, UC |
| rs10061936 | T/C | *ERAP2* | CD28 | naive | IRFs | -50 | CD, UC |
| rs9392504 | G/A | *IRF4* | TCR | naive, memory | IRFs | 93 | CeD, RA |
| rs4679081 | T/C | *TRIM71* | TCR | memory | MEF2C, IRFs | -32, -14 | CeD, MS |
| rs2115592 | T/C | *REL* | TCR | memory | IRFs, BLIMP1 | 382, 14 | CeD, RA, MS |
| 186498:20:00 | C/CT | *SOCS1* | CD28 | naive, memory | IRFs | -24 | MS |
| rs60486739 | G/A | *IL7, ZC2HC1A* | CD28 | memory | IRFs | 311 | MS |
| rs3808619 | A/C | *IL7, ZC2HC1A* | CD28 | memory | IRFs | -71 | MS |
| rs11249219 | C/T | *CLIC4* | CD28 | memory | IRFs | -24 | CeD, PSO |
| rs3784789 | C/G | *CYP1A1, CYP1A2* | CD28 | naive | SPI1 | -172 | CeD, SSc |

## Discussion

Productive T cell activation is thought to involve recognition of antigen by the TCR in association with co-stimulatory signals via receptors such as CD28 (Esensten et al., 2016). However, the relative quantities of both signals are likely to be highly variable depending on the setting of T cell activation. There is evidence to suggest that naive and memory cells have different requirements for CD28 costimulation, and a widely held view is that CD28 costimulation is less required for activation of memory than naive T cells (Croft et al., 1994; Dubey et al., 1995; London et al., 2000; Luqman and Bottomly, 1992). Understanding the requirements for CD28 costimulation during immune responses is important for many therapeutic approaches including immune suppression in autoimmunity and transplantation, as well as cancer immunotherapy.

Previous genomic studies have solely focussed on the additional effects of CD28 as a co-stimulus, and not on varying the levels of CD28 itself, and were carried out in populations of mixed cells which includes cells that have successfully undergone activation as well as the resting cells (Allison et al., 2016; Diehn et al., 2002; Wakamatsu et al., 2013). The data presented here reveal that CD28 has an important impact on memory T cells, particularly their proliferation. We were able to reach this conclusion thanks to three key components in our experimental design: (i) we specifically isolated and profiled only stimulated cells, therefore reducing the confounding effect of variable cell activation across different cell cultures, (ii) we separately assessed naive and memory cells, and (iii) we provided the first genome-wide perspective of gene expression regulation through mapping RNA and chromatin changes induced by strong CD28 stimulation in the absence of TCR. Thus, although providing CD28 on its own might not be physiological, this system allowed us for the first time to disentangle genes that are TCR or CD28 sensitive.

We observed a significant upregulation of genes in memory T cells in response to stimulation with CD28 alone that were related to cell cycle, suggesting a role for CD28 in memory cell proliferation. Indeed, we validated this directly by stimulating T cells for five days and demonstrating proliferation in memory T cells following activation with CD28 alone. Furthermore, we observed a significant upregulation of effector cytokines and chemokines in response to CD28, indicating that CD28 is also key to cell effector functions. However, our experiment was performed on a mixed memory cell population of effector and central Th cells, which differ in their levels of expression of CD28 (Koch et al., 2008). The differences in the expression levels of CD28 might have an impact in the immune response initiated, as influenza infected mice treated with CTLA4Ig shift their memory T cell pool to one predominantly composed of central memory cells, while influenza specific memory cells in untreated mice were predominantly effector cells (Ndejembi et al., 2006). This could explain why activated effector memory T cells produce cytokines more efficiently than central memory T cells (Barski et al., 2017). Finally, the variability in expression of the CD28 receptor between young and old is only significant in effector memory T cells (Koch et al., 2008). This suggests that the memory T cell activation is concordant with their ability to sense CD28, and to initiate proliferation as we demonstrate here. Therefore, the decrease of CD28 signal might contribute towards immunosenescence.

Our data provide a new context for the interpretation of several previous observations on CD28 function. Firstly, the ill-fated CD28 superagonist antibody (TGN1412) trial for B-cell chronic lymphocytic leukemia and rheumatoid arthritis, which tested the ability of CD28 costimulation to specifically expand and activate Tregs, revealed a powerful effector response to CD28 stimulation driven by effector memory T cells (Hünig, 2012). Secondly, recent data indicated that human Treg cells can be expanded by utilising CD28 antibodies alone (He et al., 2017). This is in line with our observations, given that Tregs predominantly consist of memory cells and that CD28 stimulation upregulates *FOXP3*. Thirdly, there is now increasing evidence using conditional deletion of CD28 in mice that memory T cell responses are dependent on CD28 stimulation (Fröhlich et al., 2016; van der Heide and Homann, 2016; Linterman et al., 2014; Ndlovu et al., 2014). Finally, we showed that a strong TCR signal alone is sufficient to induce the expression of key drivers of cell division and consequently trigger proliferation of naive T cells, but surprisingly had smaller effect on the proliferation of memory T cells. As such, we conclude that memory cells are not simply more sensitive to activation signals, but that there is a true difference in the TCR and CD28 requirement between the two cell types. Taken together, our data provides a genomic explanation for the requirement of CD28 in the activation and effector functions of memory T cell.

Lastly, the concept that CD28 is involved in the proliferation of memory T cells is intriguing in the light of recent data related to checkpoint blockade for cancer treatment. It has been suggested that PD-1 blockade, which is important to the reinvigoration of exhausted effector T cells, requires CD28 signaling (Hui et al., 2017; Kamphorst et al., 2017). Again, this aligns well with our data and supports the concept that differentiated memory T cells in tumours utilise CD28 and that CTLA-4 and the PD1 blockade are known to trigger autoimmunity (Ben Nasr et al., 2017; Bertrand et al., 2015).

Our findings have further implications for understanding susceptibility to complex immune-mediated diseases, where T cell activation is one of the hallmark pathobiological processes. GWAS of immune diseases have mapped hundreds of associated risk loci, many of which harbour genes of immune function. However, the specific role of the identified genes in T cell activation processes is unclear. By examining T cell gene expression sensitivity in response to specific stimuli we demonstrated that GWAS loci are enriched for CD28-sensitive genes, rather than TCR-sensitive genes, thereby increasing support for the role of T cell activation via CD28 costimulation in susceptibility of immune-mediated diseases. For example, a recent study identified that cytokine oncostatin M (OSM) is expressed at higher levels in inflamed intestinal tissues from IBD patients compared to healthy controls (West et al., 2017). In our dataset *OSM* is CD28-sensitive and is among the genes driving the enrichment of CD28-sensitive genes in IBD. The importance of CD28 costimulation in immune-mediated diseases is further supported by data from the CTLA-4 field (Walker and Sansom, 2015). Indeed, loss of CTLA-4 in mice and heterozygous mutations in humans reveal profound autoimmunity where enteropathy is a consistent feature (Kuehn et al., 2014; Schubert et al., 2014; Tivol et al., 1995). The fact that the CTLA-4 pathway directly regulates CD28 stimulation by competing for the same ligands strongly suggests that CD28 plays a key role in susceptibility to immune-mediated diseases and that increased CD28 costimulation may be sufficient for effector T cells to support their survival, proliferation and secretion of proinflammatory cytokines (Khattri et al., 1999; Qureshi et al., 2011). Our data shows that memory T cells are highly sensitive to CD28 stimulation, which is consistent with these cells being under the control of CTLA-4.

Taken together, our study provides new insights into the role of TCR and CD28 costimulation in the activation and proliferation of human naive and memory CD4 T cells, and the influence of these stimuli on immune disease susceptibility.

## Acknowledgements

We thank Sarah Teichmann for discussions and feedback. We thank Harm-Jan Westra, Gillian Griffiths, Sophie Hambleton, Neil Halliday and Roser Vento for critical feedback on the manuscript. **Funding**: We thank the Wellcome Trust for their funding supporting this research (GT, grant WT206194; DMS, grant WT204798). LJ is supported by the Wellcome Trust and the Royal Society (208750/Z/17/Z), and the Kennedy Trust for Rheumatology Research. **Authors contributions**: DAG and BS conceived and designed the project, carried out the experimental work and wrote the manuscript. DAG performed the genomic data analysis. BS carried out the FACS analysis and cell proliferation assays. LJ provided feedback on the analysis and contributed to writing the manuscript; DMS and GT conceived and designed the project, supervised the analysis, and wrote the manuscript; **Competing interests**: The authors have no conflicting of interests. **Data availability**: Upon acceptance we will make the data publically available through the European Genome-Phenome Archive (EGA; http://ega-archive.org). All the codes used in the data analysis will be made available through github upon manuscript acceptance.

## Methods

### Sample collection and DNA isolation

Blood samples were obtained from eight healthy adults, aged from 22 to 46 years. Peripheral blood mononuclear cells (PBMCs) were isolated using Ficoll-Paque PLUS (GE healthcare, Buckingham, UK) density gradient centrifugation. CD4+ T cells were isolated from PBMCs using EasySep® CD4+ enrichment kit (StemCell Technologies, Meylan, France) according to the manufacturer’s instructions. DNA was isolated from live PBMCs using Qiagen DNeasy blood and tissue kit.

All samples were obtained in accordance with commercial vendor’s approved institutional review board protocols and their research use was approved by the Research Ethics Committee (reference number: 15/NW/0282).

### Flow cytometry and cell sorting

CD4+ enriched cells were stained with the following antibodies for sorting: CD4 (OKT4)-APC (Biolegend) (AB_571944); CD25 (M-A251)-PE (Biolegend) (AB_2561861); CD127 (eBioRDR5)-FITC (eBioscience) (AB_1907343) and Live/Dead fixable blue dead cell stain. Live conventional T cells (Tcons, CD4+ CD25low CD127high) were isolated and cultured for 16h. Stimulated naive and memory cells were sorted based on the expression of CD25-PE, CD45RA-Alexa700 (Biolegend) (AB_493763) and DAPI. Resting naive and memory cells we sorted for low expression of CD25 (proportion of CD25+ cells < 1%, see **Suppl. Fig S1**).

### Cell Culture and stimulation

Chinese hamster ovary (CHO) cells expressing CD86 or FcR (FcRγII, CD32) were cultured in DMEM (Life Technologies, Paisley, UK) supplemented with 10% v/v FBS (Sigma, Gillingham, UK), 50 U/ml penicillin and streptomycin (Life Technologies), and 200 μM l-glutamine (Life Technologies) and incubated at 37°C in a humidified atmosphere of 5% CO2. CHO cells expressing CD86 and FcR were generated as previously described (Qureshi et al., 2011). T cells were co-cultured with glutaraldehyde fixed CHO-CD86 to provide CD28 signal or CHO-FcR for 16 hours in RPMI 1640 supplemented with 10% v/v FBS, 50 U/ml penicillin and streptomycin, and 200 μM l-glutamine. Cultures stimulated with CHO-CD86 were treated with with anti-CD3 (clone OKT3). Here high TCR corresponds to 1 ug/ml, low TCR corresponds to 0.01 ug/ml; high CD28 corresponds to 1:2.5 CHO-CD86 to T cell ratio and low CD28 corresponds to 1:25 CHO-CD86 to T cell ratio. Cultures stimulated with CHO-FcR were treated with 1 μg/ml of anti-CD3 (OKT3) or 1 μg/ml of anti-CD28 (9.3).

### T cell proliferation assay

Prior to stimulation with CHO-FcR cells and anti-CD3 or anti-CD28, naive and memory T cells were labeled with CellTrace Violet dye (Life Technologies) according to the manufacturer’s instructions. Five days following stimulation, T cell proliferation was analyzed by flow cytometry and proliferation was modeled using the Flowjo proliferation platform. Total T cell numbers per sample were established relative to AccuCheck counting beads (Invitrogen).

### RNA-seq

Naive and memory T cells were placed in 0.5ml of Trizol (Invitrogen) and stored at -80°C. Samples were thawed at 37°C before adding 100 μl chloroform. After reaching equilibrium, samples were centrifuged for 15 min at 4°C at 10,000g. The collected aqueous phase was mixed 1:1 with 70% ethanol before proceeding with minElute columns (Qiagen) for purification, according to the manufacturer’s protocol. RNA was quantified using Bioanalyzer. The RNA was sequenced in two separate batches. Libraries were prepared using Illumina TruSeq index tags and sequenced on the Illumina HiSeq 2500 platform using V4 chemistry and standard 75 bp paired-end. The first batch consisted of 56 samples that were multiplexed at equimolar concentrations and sequenced across 14 lanes, to yield on average 71.3 million reads per sample. The second batch consisted of 18 samples that were multiplexed at equimolar concentrations and sequenced across 3 lanes to yield on average 61 million reads per sample.

### RNA-seq data processing

Sequence reads were aligned to the GRCh38 human reference genome using STAR (v2.5.0c) (Dobin et al., 2013; Qureshi et al., 2011) and the Ensembl reference transcriptome (v83). Gene counts were estimated using featureCounts (v1.5.1) tools (Liao et al., 2014) from the subread package and only reads assigned to the transcripts were used for further processing (84-90% of reads were assigned).

To find genes that were upregulated upon stimulation, we first defined differentially expressed genes by performing pairwise comparison of all the conditions to the resting state, in a cell type specific manner, using Benjamini-Hochberg controlled FDR of 5% and an absolute fold-change ≥ 2.

We then build a linear and a switch model of gene expression using the LRT algorithm of DESeq2 (v1.14.1) separately for naive and memory cells. In the linear model, we assumed a linear increase of gene expression along with stimulus intensity (incremental fold-change ≥ 1.5 in gene expression). Genes that did not follow the linear model were tested for the switch model. Here, we assumed an “on-and-off” mode of expression, where a gene is significantly upregulated (fold-change ≥ 2) in response to the presence of either CD28 or TCR (**Fig. 1B**). In both of these models, we used all seven conditions, e.g. when testing for CD28-sensitive genes we grouped the TCR alone stimulation with the resting, since neither received CD28 signal. A gene was classified in one of the two categories without overlap and prioritised for the linear model.

To control for the different batches in which we processed the blood, which accounted for 12% of the observed variability, we performed batch correction prior to PCA using the combat algorithm as implemented by the sva package (Leek et al., 2012). To estimate the percentage of the variance explained separately by each of the recorded variables, such as the stimulus, the cell type, the gender of the donors etc., we fitted a linear model with only the stimulus or the cell type as variables (method adapted from (McCarthy et al., 2017)).

### Pathway enrichment

We performed pathway enrichment analysis by testing whether different gene-sets were over-represented in particular hallmark pathways (Leek et al., 2012; Liberzon et al., 2015). We used the Jaccard index to quantify the proportion of stimuli-specific genes present in a tested pathway and assessed the statistical significance of the over-representation using a permutation strategy. For that, within each cell type and condition, we randomly sampled 10,000 gene sets of the same size as in the observed dataset, matching for the gene size and gene expression levels.

### ChIPmentation-seq (ChM-seq)

ChIPmentation-seq was performed according to published protocol (Schmidl et al., 2015), with the following modifications to make it compatible with the iDeal Histone ChIP kit (Diagenode) buffers. Five hundred thousand crosslinked cells were washed using 250μl IL1 buffer and resuspended in 250μl IL2 lysis buffer, both of which contained 1× protease inhibitor cocktail (PIC). Cells were left to lyse for 5 minutes at 4°C on a rotator and then centrifuged at 4°C (3000g) for 5 minutes. The pellets were resuspended in 250μl IS1 buffer and sonicated using the Bioruptor®Pico (Diagenode, Belgium) for 5 minutes for resting cells or 4 minutes for stimulated cells (Diagenode). We kept a portion of the chromatin from two samples aside, one naive and one memory stimulated with highTCR and highCD28, and used them as a ChM-seq input.

The chromatin was immunoprecipitated using protein-A coated IP beads. Twenty microliters of beads were washed four times using 40μl IC1 buffer on the magnetic rack before being resuspended in 20μl of IC1. The beads were mixed with 56μl 5× IC1 buffer, 6μl 50X BSA, 1.5μl 200X PIC and 1ug antibody. We added to the mix 100μl of chromatin (equivalent to 200,000 cells) and incubated the samples overnight at 4°C at 10 rpm.

The beads were then washed on the magnet with 350μl of iW1, iW2 and iW3 buffers and a final wash with 2X 1000μl 10 mM Tris pH 8. The beads with the chromatin, as well as the two input samples, were then resuspended in 29μl ChM buffer (Tris pH 8 1M, MgCl2 1M, ChIP grade water) with 1μl Tn5 and incubated for 10 minutes at 37°C at 1500 rpm. The tagmentation was stopped with the addition of 2X 350μl iW3. Finally, the beads were washed with 350μl iW4. The chromatin from the beads was eluted using 96μl iE1 and incubated for 30min at room temperature at 1500 rpm. The chromatin was reverse cross-linked overnight using 4μl of iE2 buffer. The DNA was then purified using the MinElute PCR CLeanup kit (QIAGEN) and eluted in 30μl of water. Sequencing libraries were prepared using Nextera primers as described in the ATAC-seq protocol (Buenrostro et al., 2013). Eighteen libraries were indexed and pooled in equimolar concentration and sequenced on three lanes using the Illumina HiSeq 2500 platform and V4 chemistry using standard 75 bp paired-end reads to yield on average 80 million reads per sample.

### ATAC-seq

ATAC-seq was performed according to published protocol (Buenrostro et al., 2013), with the following modifications. Fifty thousand cells were washed with 1ml of ice-cold PBS. The cells were then resuspended in the tagmentation buffer containing Tn5 transposase (Illumina Nextera) and 0.01% digitonin and incubated for 30 minutes at 37°C before purifying the DNA using the MinElute PCR purification kit (QIAGEN). Sequencing libraries were prepared using Nextera primers as described in the ATAC-seq protocol (Buenrostro et al., 2013). Sixteen libraries were indexed and pooled in equimolar concentration and sequenced on three lanes using the Illumina HiSeq 2500 platform and V4 chemistry using standard 75 bp paired-end reads to yield on average 65 million reads per sample.

### ChM and ATAC Data Processing

The quality of the sequence reads was assessed using the fastx toolkit and the adaptors were trimmed using skewer (v0.2.2) (Jiang et al., 2014). Reads were mapped to the human genome reference GRCh38 using the bwa mem algorithm (v0.7.9a) (Li and Durbin, 2010). We only kept uniquely mapped reads, removed PCR duplicated reads and for the ATAC we excluded mitochondrial reads using samtools (v0.1.9) (Li and Durbin, 2010; Li et al., 2009). We retained 83.3% of ATAC and 73.8% of ChM reads. Genome browser tracks were created using BEDTools (v2.22.0) (Quinlan and Hall, 2010) and the UCSC binary utilities. Furthermore, we generated insert size distributions using PICARD tools (v2.6.0) CollectInsertSizeMetrics function which can be indicative of over-sonicated chromatin and excess of adapters in the data. The mapped reads were converted into bed files and chimeras were removed. Peaks were called using MACS2 (v2.1.1) (Zhang et al., 2008) setting the parameters to -q 0.05 --nomodel --extsize 200 --shift -100 for ATAC, and –broad --broad-cutoff 0.1 --nomodel --extsize 146 for H3K27ac ChM. For ChM, all samples were downsampled to to the same read number (21.6 million reads) prior to peak calling against the input.

We used the fraction of reads in peaks (FRiP), the proportion of peaks with signal value (fold-change compared to the background or the input) greater than 10, the insert size distribution and the genome tracks to investigate the quality of our data. The median FRiP score for ATAC was 59.2% and 73.5% for H3K27ac ChM. The proportion of peaks with fold-change > 10 was 22.9% for ATAC and 4.8% for ChM. We therefore concluded that the quality of our data was high. We merged all the ATAC samples and called peaks again using the parameters described above. For the H3K27ac ChM samples, we first merged the donors within each cell type and condition and then randomly sampled 17 million reads from each into one sample to reach the same read number as in the input. Since we merged already QCed samples we used the --keep-dup flag when calling peaks with MACS2, as the PCR duplicated reads for individual samples were already removed and we expected to observe a small proportion of the same reads present in independent samples by chance. We also increased the -q value threshold to 0.1 for both assays. The resulting peak files were used as a reference to count the number of reads falling into peak regions using featureCounts, therefore generating a quantitative table of read counts specifically present across the different conditions and cell types.

To ensure our analysis focused on high confidence peaks we removed the bottom 10th percentile of peaks with the lowest read counts in each dataset and obtained a final count of 142,306 and 49,638 peaks for ATAC and ChM, accordingly. To define differentially accessible regions (DARs) and differentially modified histone regions (DMHRs) the dataset was processed using DESeq2. To find regions that were upregulated upon stimulation, we compared all conditions to the resting state and used Benjamini-Hochberg controlled FDR of 5% and an absolute fold-change ≥ 2.

### Binding Expression Target Analysis

To identify regions of the genome that changed upon stimulation, which could subsequently regulate gene expression, we used Binding Expression Target Analysis (BETA) in the plus mode (Wang et al., 2013; Zhang et al., 2008). We carried the analysis using both DMHRs and DARs, as well as the list of regions that required one or the other stimulus. For example, TCR specific DMHRs or DARs were defined as the regions that are present in TCR stimulation but not in CD28 stimulation. We used the differential gene expression output from the pairwise comparison between stimulated and resting states. The median distance of interaction between an enhancer and gene promoter is estimated at 150kb (Mumbach et al., 2017), we therefore used the same window size around the transcription start site (TSS) of the differentially expressed genes to define boundaries for the analysis, along with the transcription activation domains (TADs) identified in CD4 T cells (Javierre et al., 2016). That is, if the extended 150kb region fell beyond the TAD boundary we consider the TAD coordinates as the boundary for testing predictive effects of DMHRs and DARs on gene expression. We relied on the ATAC-seq output for the transcription factor enrichment analysis (p-value ≤ 0.01) as it generates narrow peaks allowing for a more accurate estimation of the TFBS.

### Disease SNP enrichment for stimulus-sensitive genes

We collected the GWAS data for 8 common traits from the studies listed in **Suppl. Table 4**, excluding all variants that fell within the MHC locus and using a genome wide p-value threshold of < 5×10^−8^. We defined the disease loci by mapping all the SNPs in linkage disequilibrium (LD) with the reported index SNP, using r^2^ > 0.8 calculated across the European populations present in the 1000 Genomes Project data, and extending the LD boundaries by 150kb on each side, to account for the possibility of distant gene expression regulation between enhancers and gene promoters. This resulted in 234 unique regions associated to one of the eight tested traits.

We then tested whether the stimulus sensitive genes that were defined in the linear and the switch models fell within the SNP loci boundaries more often than expected by chance using a permutation strategy. To build our null distribution we selected the same number of genes, matching for gene size and expression level. We repeated the process 10,000 times.

We tested whether any of the SNPs used to define the LD boundaries overlapped with an H3K27ac or an ATAC peak identified in the specific stimulatory condition; strong CD28 alone stimulation for CD28 sensitive genes and strong TCR stimulation for TCR sensitive genes.

The disruption of TFBS by SNPs was assessed using the SNP2TFBS database (Javierre et al., 2016; Kumar et al., 2016).

## Supplementary Figures

**Supplementary Figure 1.**
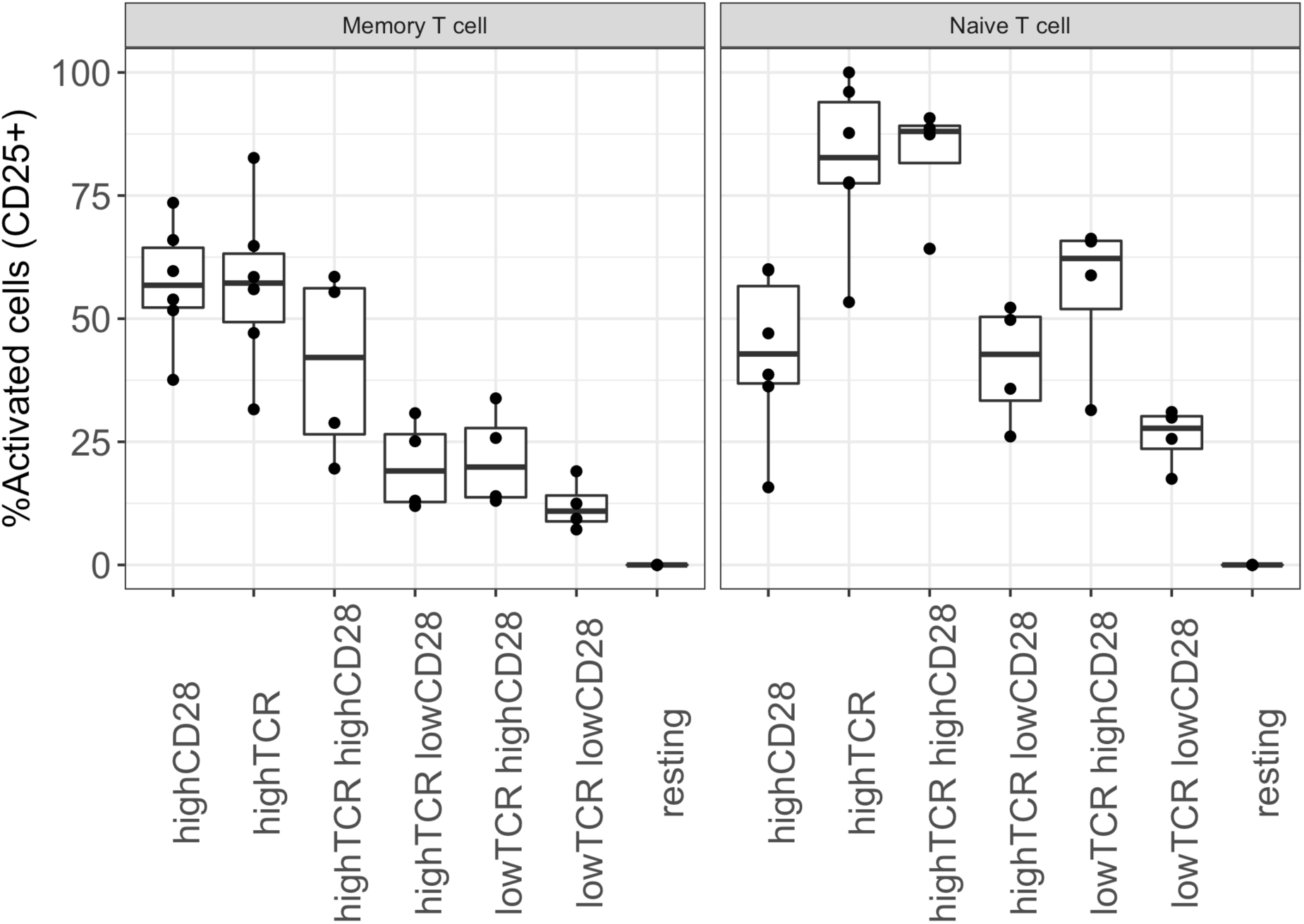
Percentage of activated naive and memory T cells upon stimulation across the experimental conditions. Cells we sorted based on the CD4+CD127+CD45RA+ for naive and CD4+CD127+CD45RA-for memory T cells. Activation was measured based on CD25 expression.

**Supplementary Figure 2.**
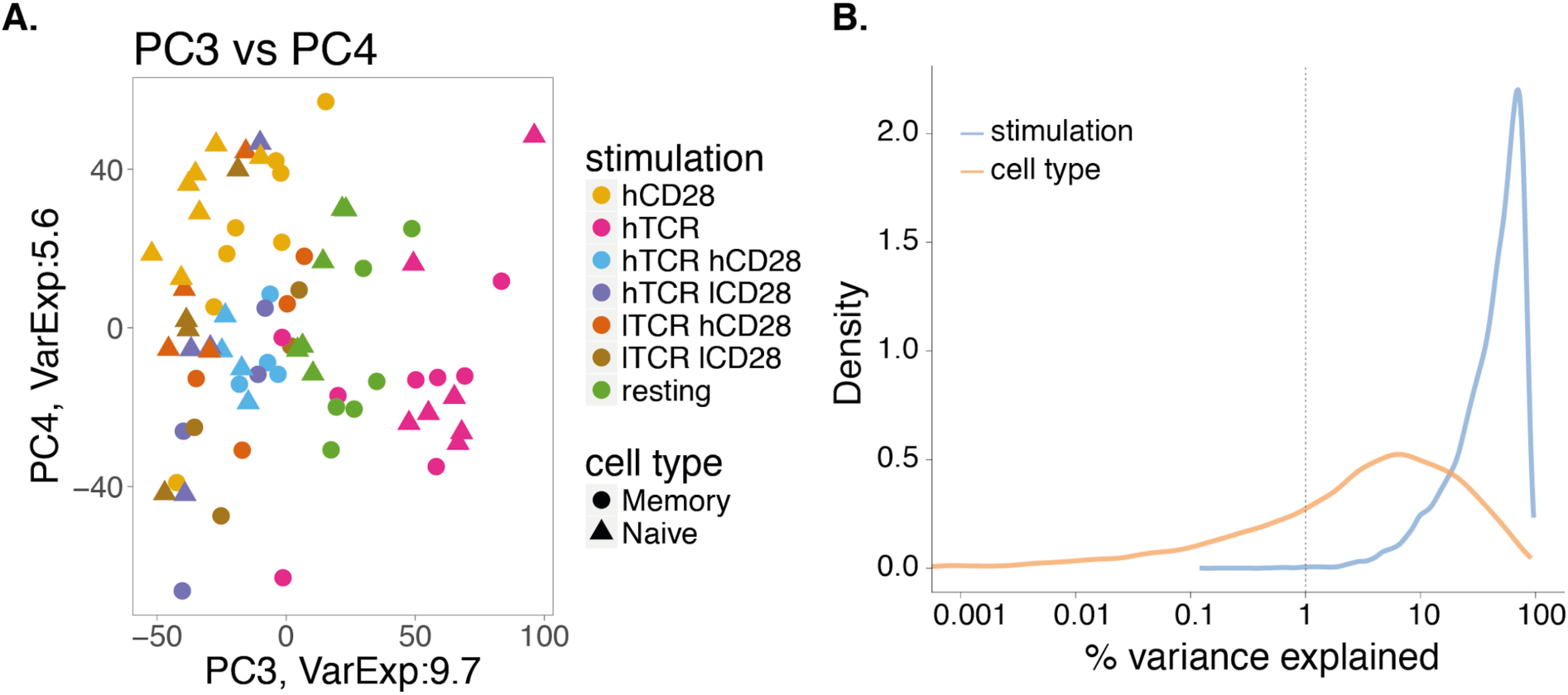
**A**. Principal component analysis using the expression of all genes. The third and fourth components explain collectively 15.3% of the observed variability and correlate with TCR and CD28 intensity. Each dot corresponds to an individual sample, colored by stimulation and shaped according to the cell type. **B**. The percentage of the total variance explained by stimulation and cell type.

**Supplementary Figure 3.**
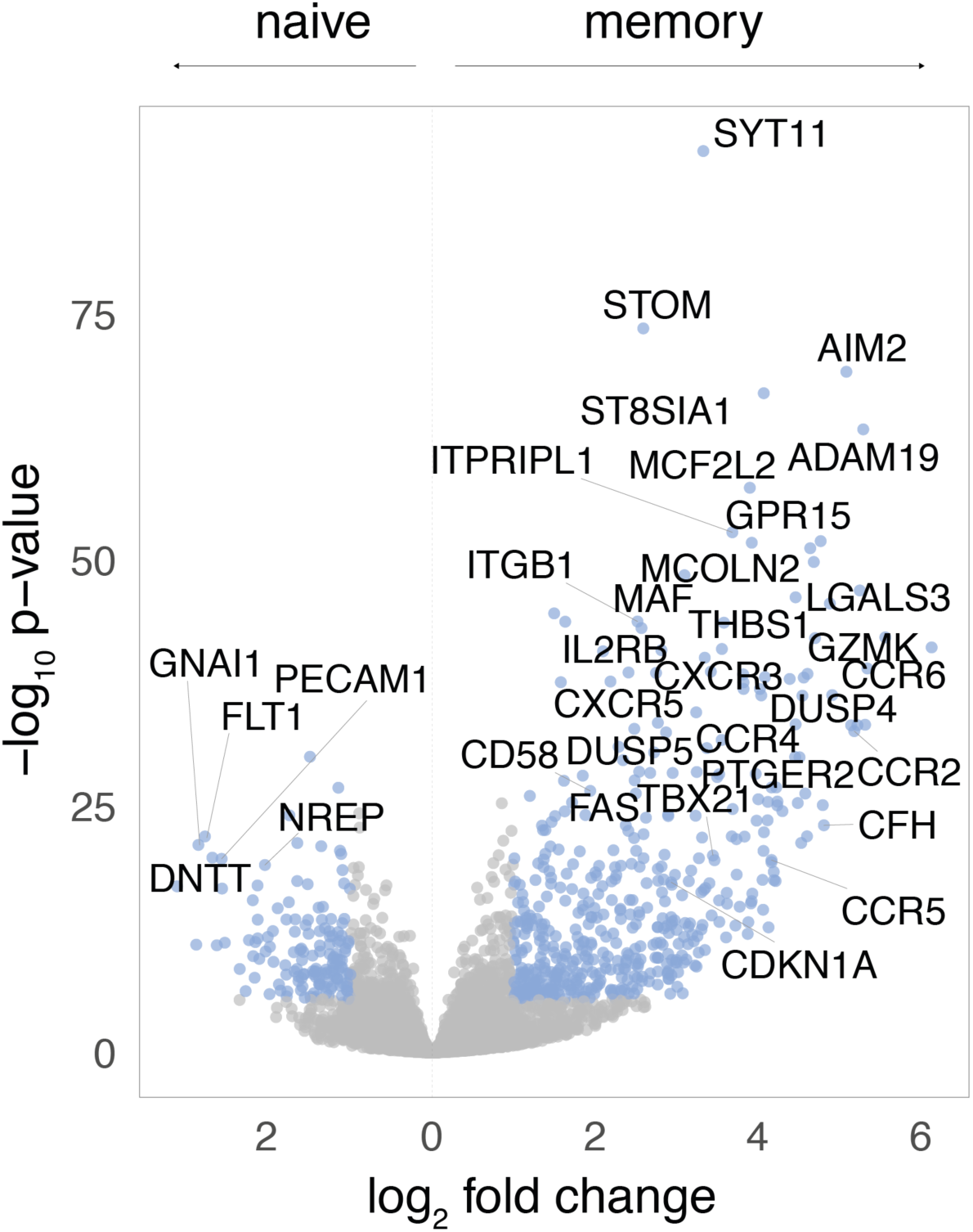
Volcano plot of differential gene expression test between resting naive cells and resting memory cells. Genes colored in blue correspond to differentially expressed genes with log2 fold-change > 1 and FDR < 5%. Labelled are DEG with the lowest p-values.

**Supplementary Figure 4.**
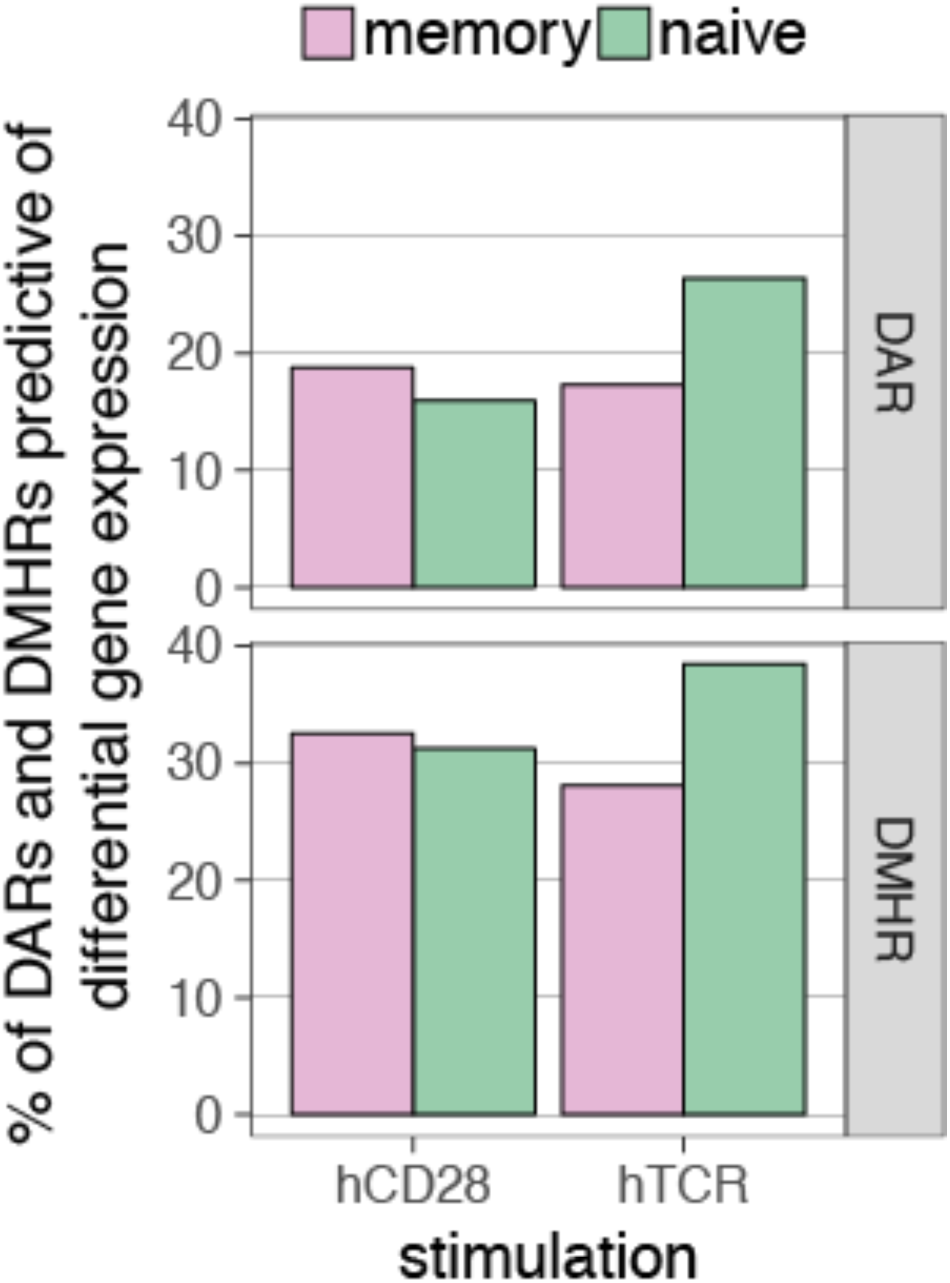
Proportion of differential accessible regions (DARs) and differentially modified histone regions (DMHRs) that regulate at least one differentially expressed gene (< 150kb away from TSS).

**Supplementary Figure 5.**
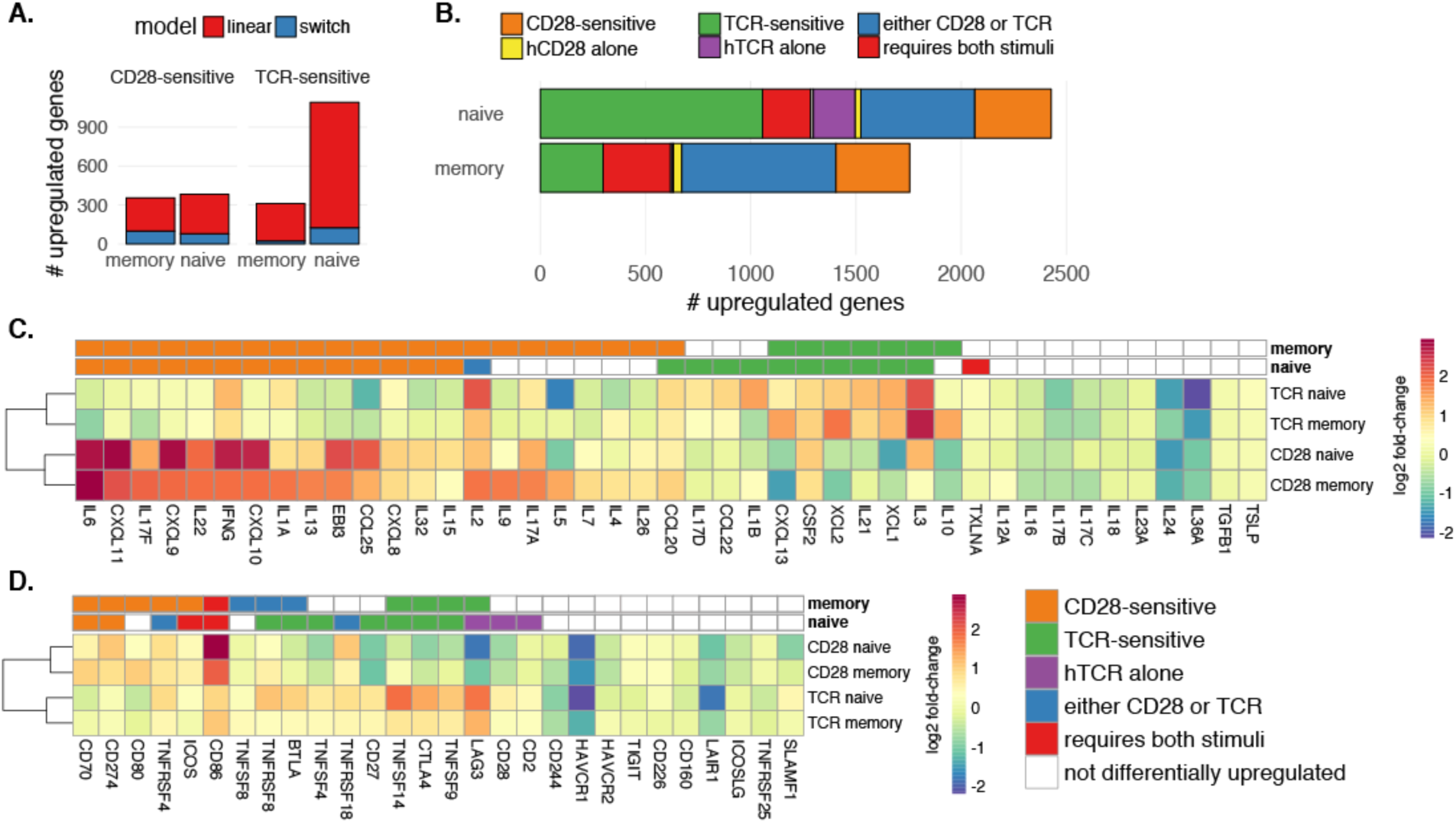
**A.** Number of TCR and CD28 sensitive genes identified by the linear (priority) and the switch model in the two cell types. **B.** Number of differentially expressed genes (DEG) upon stimulation. The coloring represents different stimulatory dependencies. Grey indicates a small number of peaks that were only shared between hTCR alone and hCD28 alone stimulations. **C+D**.Heatmap of the log2 fold-change identified using the linear model of gene expression for (**C**) cytokines and chemokines and (**D**) costimulatory ligands and receptors that are expressed in the dataset.

**Supplementary Figure 6.**
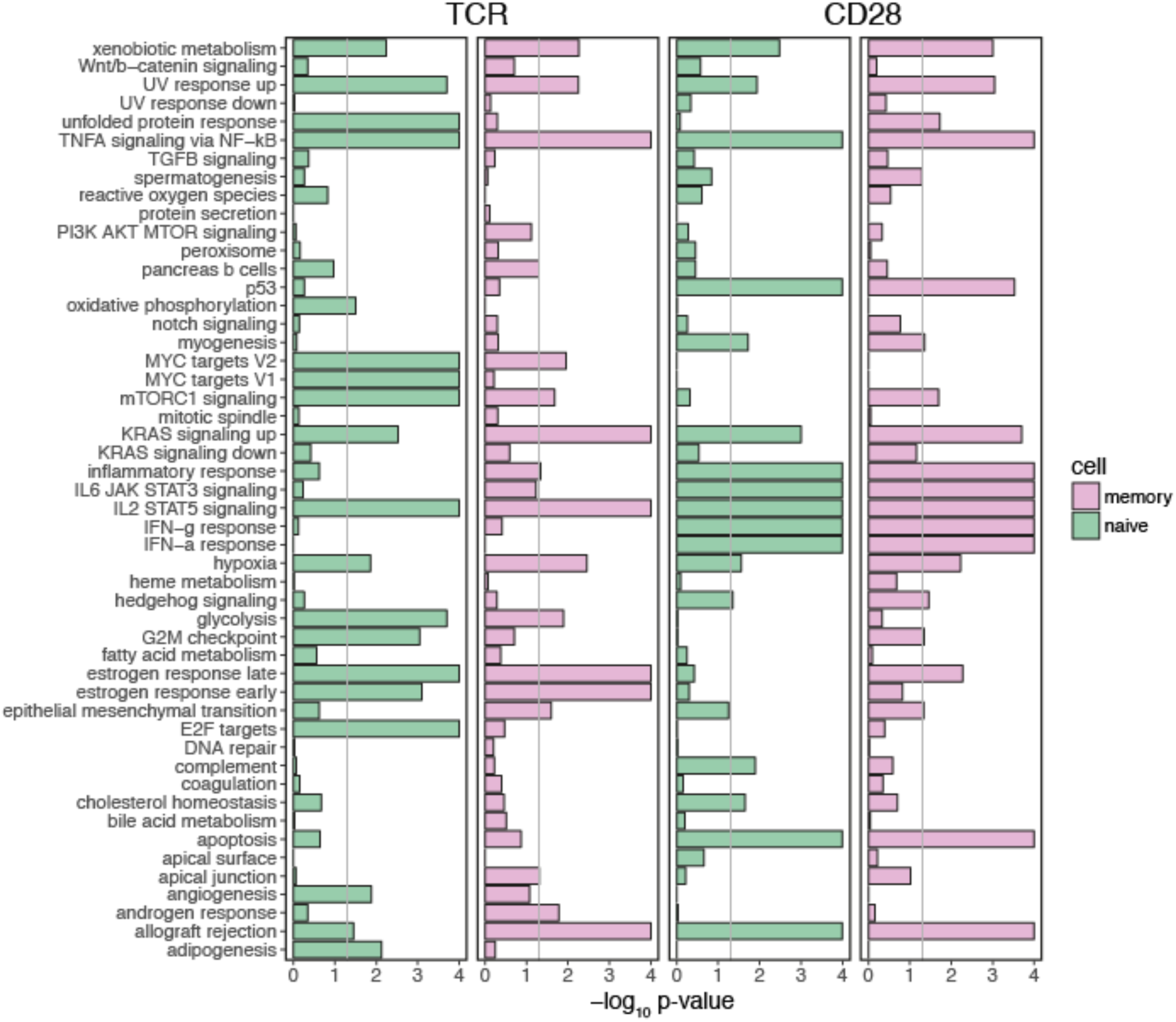
Enrichment p-values for all hallmark pathways for TCR-sensitive and CD28-sensitive genes in naive and memory cells.

## Supplementary tables

**Table S2.** Differential expression analysis output tables from DESeq2 from the RNA data linear and switch models comparisons between stimulated states and the resting state. https://drive.google.com/file/d/1qnRuVl1iCL1GfA6D5×6KHraw1QwSO59a/view?usp=sharing

**Table S1.** Differential expression analysis output tables from DESeq2 from the RNA data pairwise comparison between stimulated states and the resting state.https://drive.google.com/file/d/1QGrNAgRjVvaiHwGUnES11×0_V3SqD7g_/view?usp=sharing

**Table S3.** List of switcher genes identified in the study.

| gene | log2FC memory<br>CD28 | FDR memory<br>CD28 | log2FC naive<br>TCR | FDR naive<br>TCR |
| --- | --- | --- | --- | --- |
| *TRNP1* | 0.635 | 9.08x10<sup>-5</sup> | 0.627 | 2.06x10<sup>-3</sup> |
| *CDC20* | 0.636 | 1.56x10<sup>-4</sup> | 0.88 | 2.03x10<sup>-9</sup> |
| *AK4* | 0.756 | 0.0351 | 1.407 | 7.36x10<sup>-6</sup> |
| *DTL* | 1.153 | 8.09x10<sup>-8</sup> | 0.807 | 1.32x10<sup>-3</sup> |
| *MTHFD2* | 1.149 | 0.0134 | 0.923 | 1.04x10<sup>-4</sup> |
| *CCL20* | 1.075 | 3.34x10<sup>-4</sup> | 0.971 | 1.87x10<sup>-3</sup> |
| *TFRC* | 0.691 | 2.47x10<sup>-4</sup> | 0.91 | 5.16x10<sup>-8</sup> |
| *STC2* | 3.051 | 2.85x10<sup>-3</sup> | 1.534 | 4.81x10<sup>-4</sup> |
| *EPB41L4B* | 0.771 | 7.18x10<sup>-4</sup> | 0.968 | 3.82x10<sup>-4</sup> |
| *TXN* | 0.68 | 2.69x10<sup>-4</sup> | 0.683 | 1.86x10<sup>-4</sup> |
| *MCM10* | 1.173 | 1.62x10<sup>-8</sup> | 1.107 | 2.83x10<sup>-5</sup> |
| *DNAJC12* | 1.011 | 2.18x10<sup>-3</sup> | 1.015 | 3.72x10<sup>-5</sup> |
| *DGAT2* | *1.07* | *5.53x10<sup>-3</sup>* | 0.75 | 2.99x10<sup>-6</sup> |
| *CHEK1* | 0.727 | 2.31x10<sup>-9</sup> | 0.603 | 1.57x10<sup>-8</sup> |
| *NETO2* | 0.807 | 1.45x10<sup>-4</sup> | 1.355 | 1.59x10<sup>-5</sup> |
| *NLN* | 0.635 | 6.08x10<sup>-3</sup> | 0.919 | 5.06x10<sup>-6</sup> |
| *CDC6* | 0.658 | 1.74x10<sup>-4</sup> | 0.734 | 4.65x10<sup>-7</sup> |
| *NME1-NME2* | *1.072* | *0.033* | 1.054 | 5.67x10<sup>-7</sup> |

\* In italics are the log2 fold-change and p-values identified only in the switch model.

**Table S4.** Details of the GWA studies used in this analysis.

| **trait** | **acronym** | **source** | **GWAS_type** | **Processing notes** |
| --- | --- | --- | --- | --- |
| Bone mineral density | BMD | GWAS catalog (search term: "Bone Mineral Density") | NA | 1 |
| Crohn's disease | CD | Jostins, 2012, Nat | Meta-analysis and finemapping IBD, CD and UC | 2 |
| Ulcerative colitis | UC | Jostins, 2012, Nat | Meta-analysis and finemapping IBD, CD and UC | 2 |
| Celiac disease | CeD | Trynka, 2011 Nat Genet | ImmunoChip | 3 |
| Multiple sclerosis | MS | Beecham, 2013 Nat Genet. | ImmunoChip | 4 |
| Rheumatoid arthritis | RA | Okada, 2014 Nat | Meta-analysis GWAS and ImmunoChip | 5 |
| Psoriasis | PSO | Tsoi, 2012, Nat Genet | Meta-analysis GWAS and ImmunoChip | 4 |
| Systemic sclerosis | SSc | Bossini-Castillo 2015 Semin Immunopathol | Summary GWAS and ImmunoChip | 6 |
| Type 1 Diabetes | T1D | Onengut-Gumuscu, 2015, Nat Genet | ImmunoChip | 4 |

